# miRScore: a rapid and precise microRNA validation tool

**DOI:** 10.1101/2024.12.12.628184

**Authors:** Allison Vanek, Sam Griffiths-Jones, Blake C. Meyers, Saima Shahid, Michael J. Axtell

## Abstract

MicroRNAs (miRNAs) are small non-protein-coding RNAs that regulate gene expression in many eukaryotes. Next-generation sequencing of small RNAs (small RNA-seq) has accelerated discovery and annotation of novel miRNAs. Newly discovered miRNAs are typically submitted to databases such as the miRBase microRNA registry following the publication of a peer-reviewed study. However, genome-wide scans using small RNA-seq data often yield high rates of false-positive miRNA annotations, highlighting the need for more robust validation methods. miRScore was developed as an independent and efficient tool for evaluating miRNA annotations using sRNA-seq data. miRScore combines structural and expression-based analyses to provide rapid and reliable validation of miRNA annotations. By providing users with detailed metrics and visualization, miRScore enhances the ability to assess confidence in novel and existing miRNA annotations. miRScore has the potential to advance the overall quality of miRNA annotations by improving accuracy of new submissions to miRNA databases and serving as a resource for re-evaluating existing annotations.

## Introduction

MicroRNAs (miRNAs) are a class of small, non-coding RNAs that regulate gene expression within eukaryotes. This regulation typically occurs when a miRNA, which is loaded into an RNA-induced silencing complex (RISC), imperfectly base pairs to a target messenger RNA (mRNA). The RISC then frequently acts as an endonuclease to cleave the mRNA or to otherwise inhibit its translation (1–4). miRNA-directed regulation of mRNAs is crucial in various biological processes such as developmental timing (5–7), metabolism (8,9), and defensive pathways (10–13) in both plants and animals. Although miRNA biogenesis varies somewhat between animals and plants, the fundamental aspects of miRNA structure and function are conserved (14). In both plants and animals, the precursors of miRNAs are generally transcribed by RNA polymerase II from an endogenous *MIRNA* gene. While many *MIRNA* primary transcripts are transcribed as independent genes from intergenic regions, some are processed from the introns of protein-coding mRNAs (4). Transcription results in a long RNA, containing a hairpin, called the primary miRNA. The hairpin embedded within the primary transcript is then processed by sequential endonuclease activity to release a double-stranded miRNA duplex. The miRNA duplex is a double-stranded RNA which consists of the mature functional strand (miRNA) and passenger strand (miRNA*). The miRNA duplex is unwound and the single-stranded mature miRNA is bound to an Argonaute protein to form the RISC, while the miRNA* is typically degraded. For details of microRNA biogenesis, see (15,16,4).

Alignment of deep small RNA-sequencing (sRNA-seq) data to a reference genome is a common method for *MIRNA* annotation and quantification. Several tools such as ShortStack (17,18), miRador (19), miRDeep (20), and miRDeep-P2 (21), have been developed to annotate miRNAs and other small RNAs using sRNA-seq data. These tools typically work by aligning sRNA-seq data to a reference genome, followed by evaluation of potential miRNA-encoding loci (*MIRNA)*. One way candidate *MIRNA* loci are identified is by the distinctive alignment pattern of the miRNA duplex reads to the hairpin precursor. miRNA and miRNA* reads from sRNA-seq align to a single genomic strand, as their precursors are single-stranded transcripts. These reads align a short distance from each other, forming two distinct “stacks” of read coverage (17,22). *MIRNA* primary transcripts are short-lived and typically not detected using sRNA-seq or regular mRNA-seq. Most sRNA-seq centered *MIRNA* identification tools thus annotate “hairpin” sequences that encompass the stem-loop region and some adjacent sequence of pre-determined length. The start and stop positions of these annotations do not necessarily correspond to the actual primary transcripts. The secondary structure of this putative hairpin precursor is then predicted. For true *MIRNA* loci, the predicted secondary structure of the putative precursor RNA is an imperfect stem-loop. Furthermore, two stacks of aligned sRNA-seq reads from the miRNA and the miRNA* are found on opposite arms of the predicted stem-loop with a diagnostic two nucleotide 3’-overhang. Identification of candidate *MIRNA* loci using sRNA-seq is therefore dependent on empirical evaluation of read alignment patterns in the context of the presumed precursor’s predicted RNA secondary structure.

The identification of *MIRNAs* through deep sequencing data poses some challenges. One is the handling of multimapping reads, in which there are multiple best-scoring alignments for a single read. This occurs frequently with sRNA-seq data due to shorter read lengths and the fact that identical miRNAs can be encoded by paralogous *MIRNA* loci (18). Another challenge is distinguishing true *MIRNA* loci from other sRNA types such as short-interfering RNAs (siRNAs), which have their own unique alignment patterns and criteria (23,24). Each *MIRNA* discovery tool employs unique methods for handling these challenges, with varying degrees of performance for identification of novel *MIRNAs* in plants and animals (20,17,19,21). The lack of uniform implementation of well-defined *MIRNA* criteria, coupled with the challenging nature of informatically distinguishing miRNAs from noise or other sRNA species, has led to diminishing confidence in the overall quality of existing miRNA annotations (25). There have been considerable efforts to define miRNA criteria to improve the quality of annotations (23). However, databases such as miRBase rely on researchers and peer reviewers to assess the validity of miRNAs before submission, and have adopted methods of assessing confidence rather than gatekeeping submitted annotations (25,26). In addition to the quality of annotations, the submission and assessment of novel miRNAs relies on manual inspection and evaluation. This process significantly delays cataloging efforts and has created a backlog of submissions, highlighting a need for an automated method of validation.

While there are a growing number of sRNA annotation tools and miRNA databases available, the community lacks a tool to quickly analyze novel and annotated *MIRNA*s for further analysis or submission to a database. Such a tool would provide the validation expected of novel submissions without manual scrutinization, reducing the time it takes to analyze and catalog miRNA annotations for both researchers and database curators. To address this need, we developed miRScore – a rapid and precise miRNA validation tool. miRScore can rapidly evaluate both existing and novel miRNAs using widely accepted *MIRNA* criteria in plants and animals. It offers a comprehensive evaluation of *MIRNA* loci, analyzing each criterion and producing visualizations of hairpin secondary structure and expression patterns. In this study, miRScore is described and tested using published and novel *MIRNAs* from plants and animals.

## Design and Implementation

miRScore is implemented as a Python script that requires several commonly used bioinformatic tools including samtools (27), ViennRNA (28), and bowtie (29). miRscore is an open-source software available under a permissive MIT license from GitHub at https://github.com/Aez35/miRScore, and is easily installed using Bioconda (30).

### Workflow

miRScore validates *MIRNA* loci by analyzing the hairpin precursor sequence, mature miRNA sequence, and sRNA-seq data. The validation process uses a set of previously described criteria which can be categorized as either structural (based on the predicted RNA secondary structure of the precursor) or expression (based on observations of miRNA and miRNA* abundance) (Table 1)(31,25,32,23).

**Table 1.**
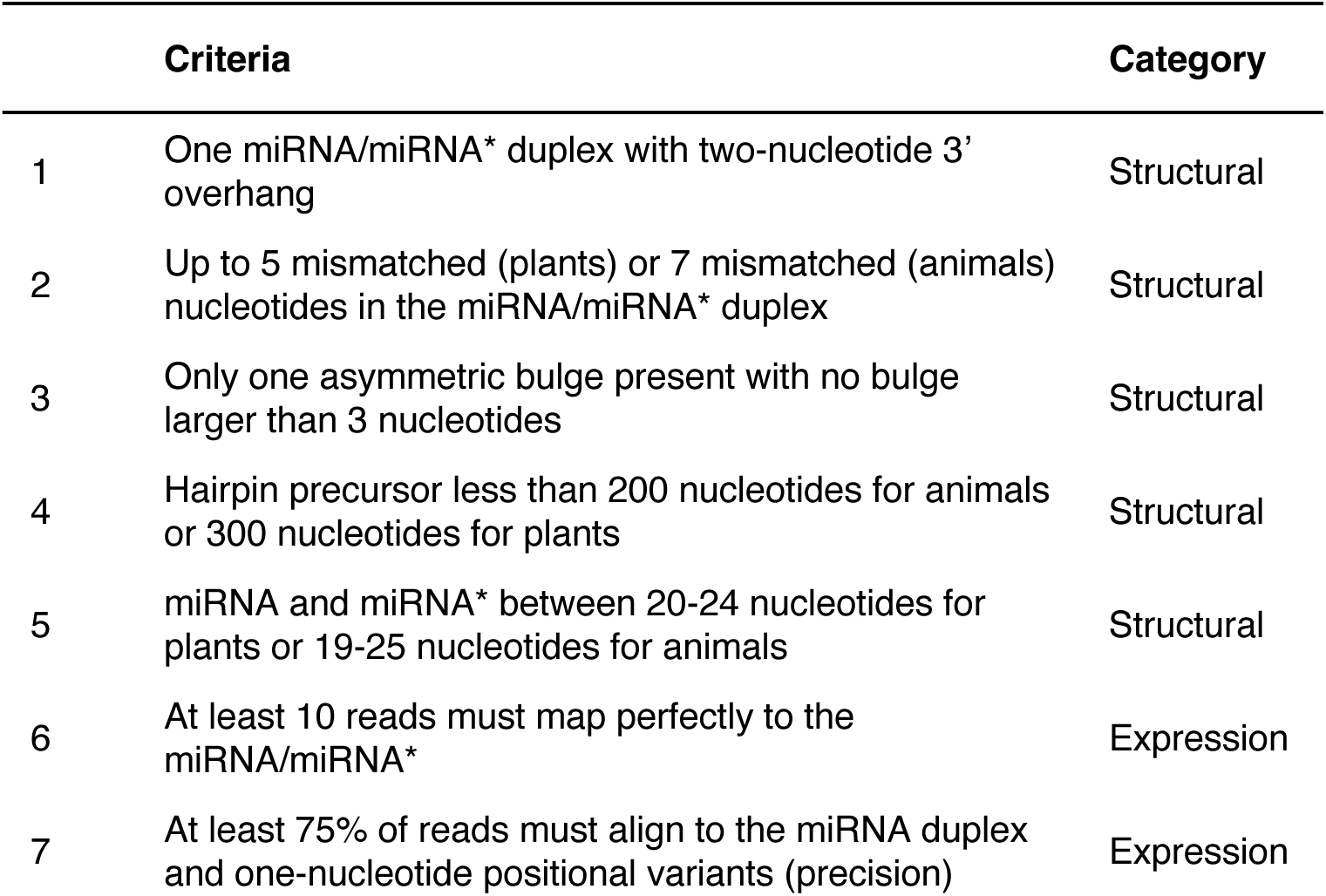
Criteria for endogenous miRNAs in plants and animals.

|  | Criteria | Category |
| --- | --- | --- |
| 1 | One miRNA/miRNA* duplex with two-nucleotide 3' overhang | Structural |
| 2 | Up to 5 mismatched (plants) or 7 mismatched (animals) nucleotides in the miRNA/miRNA* duplex | Structural |
| 3 | Only one asymmetric bulge present with no bulge larger than 3 nucleotides | Structural |
| 4 | Hairpin precursor less than 200 nucleotides for animals or 300 nucleotides for plants | Structural |
| 5 | miRNA and miRNA* between 20-24 nucleotides for plants or 19-25 nucleotides for animals | Structural |
| 6 | At least 10 reads must map perfectly to the miRNA/miRNA* | Expression |
| 7 | At least 75% of reads must align to the miRNA duplex and one-nucleotide positional variants (precision) | Expression |

miRScore utilizes a “pass/fail” system of reporting, in which a locus must meet all criteria or it fails. miRScore was envisioned to be used to assess the quality of new *MIRNAs* prior to submission to microRNA databases and for evaluating the support for existing microRNA annotations (Figure 1A). Users input properly formatted FASTQ files containing sRNA sequencing reads, mature miRNA sequences, and hairpin sequences in FASTA format (Figure 1B). The precursor sequences should be extended past the miRNA Dicer/DCL cut site, and the miRNA/miRNA* should not start or end the precursor sequence. This is to allow proper evaluation of the miRNA duplex structure. The identifier of each *MIRNA* locus should be identical to the corresponding mature miRNA. Multiple *MIRNA* loci may be submitted for a single mature miRNA. Users may include miRNA* sequences in the mature FASTA file, but they must be distinguishable from the mature sequence by the either a “-5p”,”-3p”, “.star”, or “ * “ at the end of the name (i.e. miR399-3p, miR399.star, miR399*) (Figure 1C). miRScore evaluates structural and expression criteria of all loci, assigns a pass or fail result, reanalyzes each failed locus, and generates visualizations.

**Figure 1.**
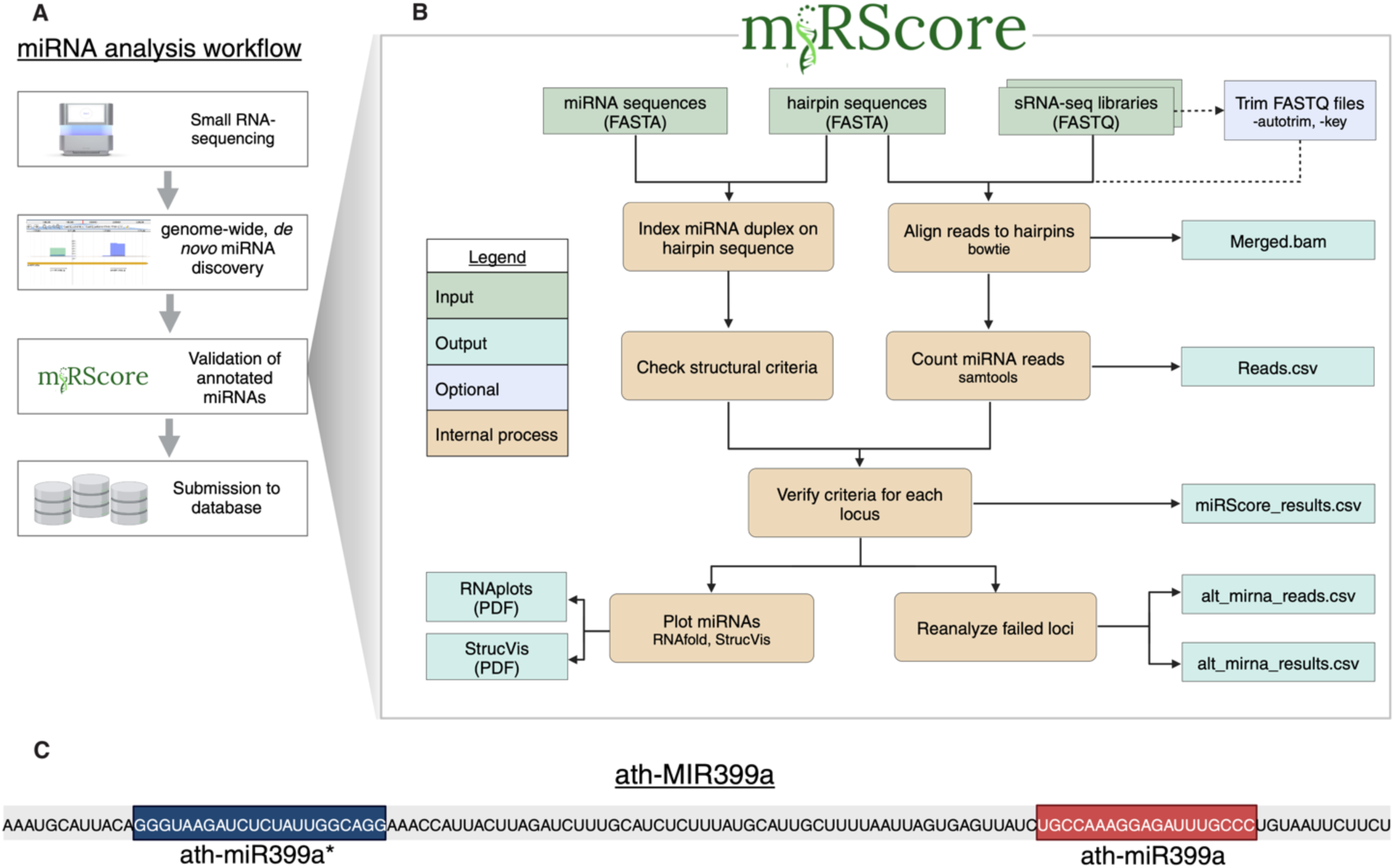
Workflow and input of miRScore. (A) miRScore is designed to follow miRNA annotation in the miRNA analysis workflow. (B) Flow chart describing the inputs and steps of miRNA analysis by miRScore (C) Example of suitable names for sequences for input FASTA files. *MIRNA* hairpin identifier (ath-MIR399a) must match the mature miRNA sequence identifier (ath-miR399a), however the miRNA* (ath-miR399a*) must have an identifier that distinguishes it from the mature miRNA sequence within the file.

### Structural evaluation

miRScore predicts the secondary structure of single-stranded hairpin precursors using RNAfold from ViennaRNA (28). The location of the miRNA and miRNA* sequences are indexed on the hairpin. If the user does not provide a miRNA* sequence, miRScore predicts it by determining the sequence that forms a miRNA/miRNA* duplex with a two-nucleotide 3’ overhang. miRScore then evaluates the miRNA duplex and hairpin against structural criteria (Table 1). This secondary structure is used to determine characteristics such as the number of mismatches and large bulges.

### Expression evaluation

The next step of the process is to evaluate expression-based criteria (Table 1). In this phase, miRScore quantifies miRNA abundance and calculates precision for each *MIRNA* locus. Reads from each library are mapped to the hairpin using bowtie (29). Alignments are output as separate bam files for each library. When counting miRNA and miRNA* reads, miRScore allows for one-nucleotide positional variance, which is included to account for biological variation in Dicer/DCL processing during miRNA biogenesis (23). At least 10 exact reads from a single library must map to the miRNA duplex. Reads from both the miRNA and miRNA* must be detected. miRScore then calculates precision for each locus. Precision is defined as the number of miRNA duplex reads divided by the total number of reads which map to the hairpin precursor. At least 75% of the reads mapping to a hairpin must map to the miRNA duplex or its one-nucleotide positional variants to meet the precision threshold.

### Identifying potential alterative mature miRNAs in failed loci

Any *MIRNA* locus that fails to meet all *MIRNA* criteria is reanalyzed to determine if a potential mature miRNA exists at the failed locus. Reanalysis begins by determining the most abundant reads, 20-24 nucleotides in length, that map to the failed hairpin sequence. miRScore then evaluates this sequence as an ‘alternative miRNA’ using structural and expression criteria (Table 1). If all criteria are met, miRScore includes this potential ‘alternative miRNA’ in a separate alternative results CSV file. Read counts for all alternative miRNAs are reported in an additional alternative reads CSV file.

### Output and visualization

After assessing structural and expression-based criteria, miRScore generates a CSV file containing details about each locus along with a pass or fail result. If a *MIRNA* locus fails any criteria miRScore will report the locus as failed. Reasons for failure are described in the ‘flags’ column of the results CSV file. Lastly, miRScore generates figures for each submitted *MIRNA* locus for visualization of secondary structure and read depth.

## Results and Discussion

### Performance analysis for annotated miRNAs

miRScore was used to examine existing *MIRNA* annotations sourced from miRBase version 22.1 across five species including: *Arabidopsis thaliana, Homo sapiens, Mus musculus, Oryza sativa,* and *Zea mays* (Table 2).

**Table 2.**
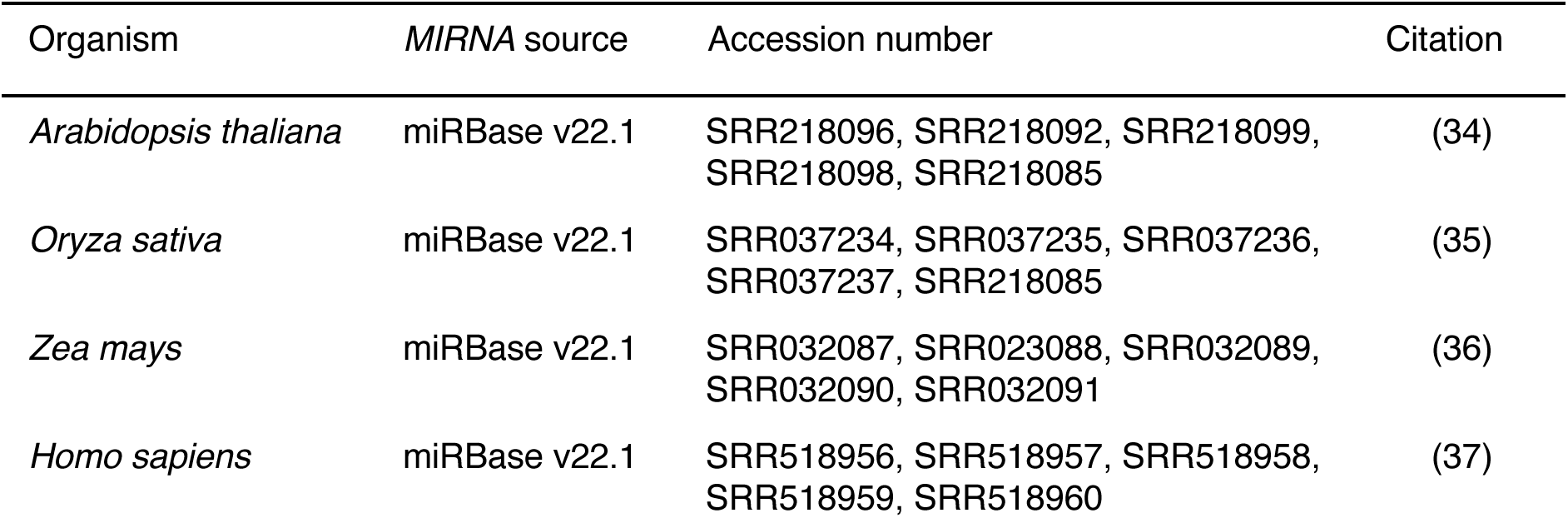
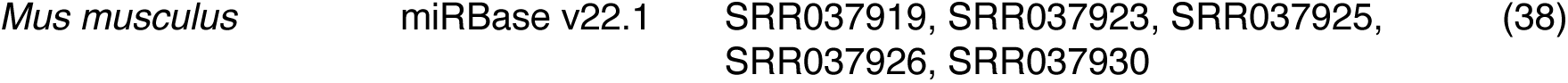
*MIRNA* and small RNA-seq data sources.

For each species, five sRNA-seq libraries were acquired from the most frequently cited publication that included suitable sRNA-sequencing data (33–37). miRScore was executed using default settings. There was a positive correlation between runtime and the number of *MIRNAs* analyzed (Figure 2A). However, there was not a positive correlation between runtime and the number of small RNA-seq reads processed (Figure 2B). These findings suggest that miRScore’s runtime is influenced more by the number of annotations examined than by the depth of small RNA sequencing data.

**Figure 2.**
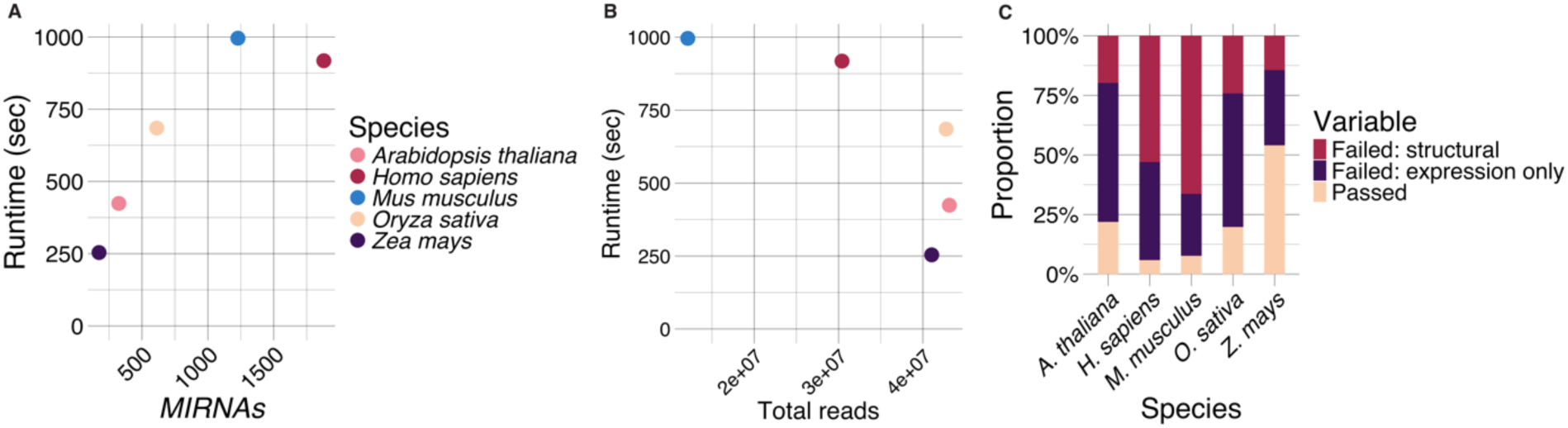
Performance of miRScore in five annotated *MIRNA* datasets. (A) Total runtime (seconds) versus number of *MIRNA*s in dataset. (B) Total runtime (seconds) versus number of reads in five sRNA-seq libraries for each dataset. (C) Proportion of *MIRNAs* with miRScore result passed (tan), failed due to structural criteria (red), or failed due to expression criteria only (purple).

The primary output of miRScore is a pass/fail result for each locus, accompanied by flags which indicate specific criteria a *MIRNA* locus did not meet (Supplemental File S1). A single *MIRNA* locus may receive multiple flags if it fails to meet multiple criteria. We evaluated the distribution of failed *MIRNA*s across structural and expression-based categories for all tested species (Figure 2C). When a locus failed due to both structural and expression criteria, it was categorized as a structural failure, as structural criteria remain consistent across analyses, whereas expression can vary depending on library input.

In animal datasets, most of the failed *MIRNA* loci failed due to structural criteria (Figure 2C). For instance, 709 out of 1770 failed *H. sapiens MIRNAs* had annotated miRNA and miRNA* sequences that did not form a miRNA duplex with a two nucleotide 3’ overhang, often missing a single nucleotide on either the 5’ or 3’ end (Supplemental Figure S1, Supplemental File S1). In contrast, failure to meet expression criteria was the most common cause of failure for plant *MIRNA*s (Figure 2C). For example, of the 256 failed *A. thaliana MIRNAs*, 168 had no detectable reads in the analyzed sRNA-seq data, and 47 had a precision of less than 75% (Supplemental Figure S1, Supplemental File S1). Significant fractions of miRBase-annotated *MIRNA* loci did not pass miRScore evaluation. Some of the failures are attributed to tissue-specific or conditional accumulation of the mature miRNA such that the miRNA and/or the miRNA* were absent in the sRNA-seq data used for analysis. Other failed loci likely reflect the subset of miRBase annotations that are not true *MIRNAs* (38,39). miRScore visualizations of hairpin secondary structure and read depth for each input locus (Figure 3A-D) allows intuitive inspection of results with respect to the *MIRNA criteria* (Figure 3E). For example, inspection of the visualizations of *ath-MIR399a* (Figure 3A-B), an endogenous *A. thaliana MIRNA*, visually confirms that this locus meets all criteria (Figure 3E). Conversely, *ath-MIR405a* (Figure 3C-D) failed miRScore analysis due to unmet expression criteria (Figure 3E).

**Figure 3.**
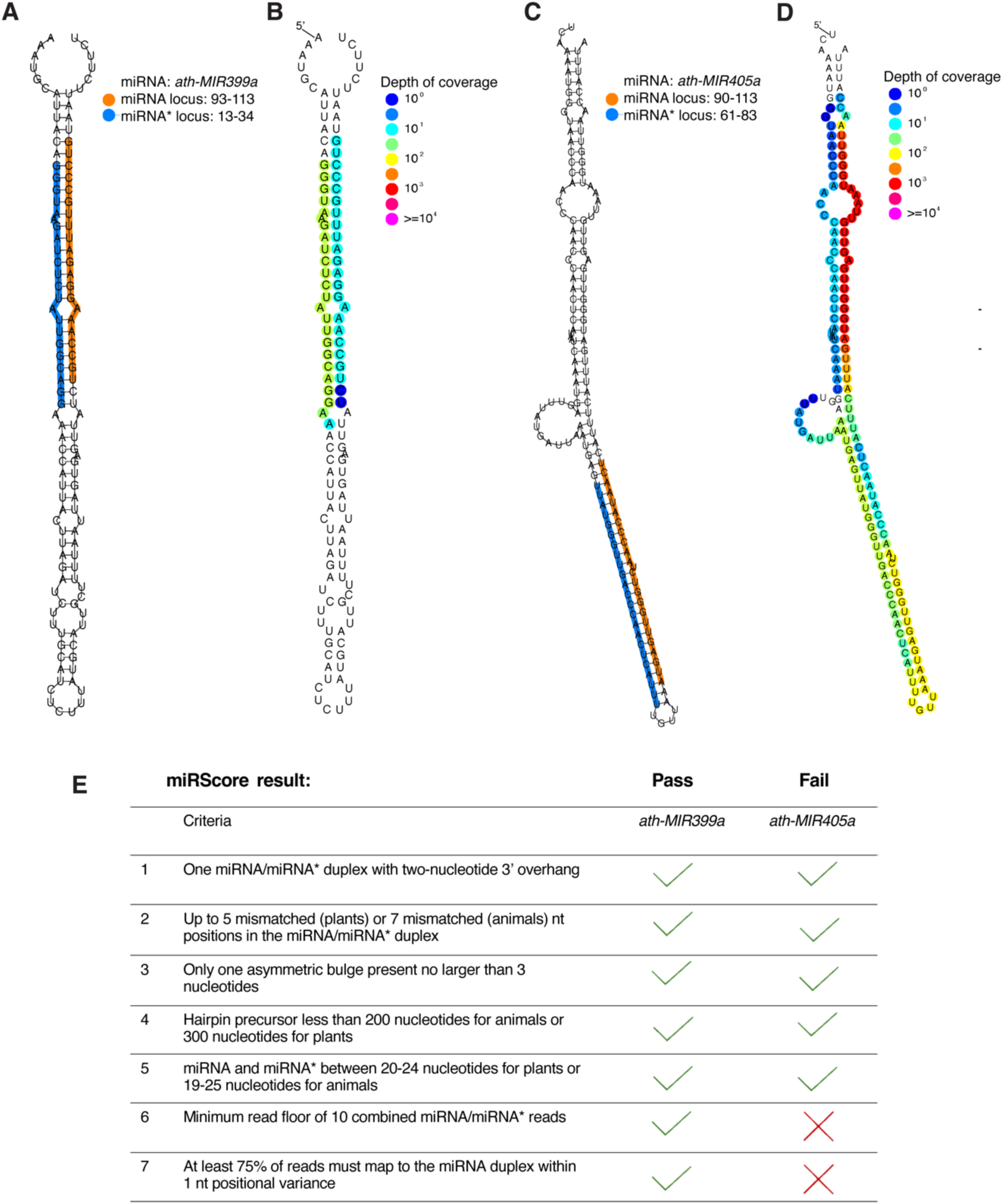
Visualization of RNA secondary structure and read depth for example *MIRNA*s. (A) *ath-MIR399a* passed all criteria. RNAplot (left) and Strucvis plot (right) of *ath-MIR399a* visually depicts many of the criteria. Mature miRNA (orange) and miRNA* (blue) in RNAplot indicates where the user-provided sequence can be found within the hairpin precursor secondary structure. Read depth from all user-submitted libraries can be seen in the Strucvis plot. (B) *ath-MIR405a* failed two criteria. RNAplot (left) and Strucvis plot (right) of *ath-MIR405a* visually depicts many of the criteria. Mature miRNA (orange) and miRNA* (blue) in RNAplot indicates where the user-provided sequence can be found within the hairpin precursor secondary structure. Read depth from all user-submitted libraries can be seen in the Strucvis plot.

One feature of miRScore designed to help users evaluate failed loci is the reanalysis step. In this final step, miRScore examines each *MIRNA* locus that failed validation and determines if there is another potential miRNA at the locus based on alignment of sRNA-seq data to the hairpin. The most abundant reads that map to the hairpin is assigned as an ‘alternative’ miRNA, which is then evaluated using structural and expression criteria. This feature rescued several *MIRNA* loci across all tested species in the dataset acquired from miRbase (Supplemental File S2).

### Manual validation of annotated MIRNAs

To evaluate miRScore’s classification performance, each *MIRNA* locus across all species was manually inspected to determine its actual condition (pass or fail). Manual inspection used a combination of data including plots of RNA secondary structure overlaid with annotation and alignment data (Figure 3A-D) and genome browser visualizations of aligned small RNA-seq data. No false positives or false negatives were observed in any of the analyzed results (Table 3).

**Table 3.** miRScore performance metrics.

| Organism | *MIRNAs* submitted to miRScore | True positive *MIRNAs* | True negative *MIRNAs* | False positive *MIRNAs* | False negative *MIRNAs* |
| --- | --- | --- | --- | --- | --- |
| *Arabidopsis thaliana* | 324 | 68 | 256 | 0 | 0 |
| *Oryza sativa* | 612 | 119 | 493 | 0 | 0 |
| *Zea mays* | 174 | 94 | 80 | 0 | 0 |
| *Homo sapiens* | 1880 | 92 | 1788 | 0 | 0 |
| *Mus musculus* | 1228 | 117 | 111 | 0 | 0 |

### Performance analysis for de novo MIRNAs

miRScore was designed to evaluate *MIRNA* annotations. One common source of annotations are those produced by tools that perform genome-wide *de novo* annotation such as ShortStack (17), miRador (19), and miRDeep-P2 (21). We generated *de novo MIRNA* annotations in four plant species: *Oryza sativa, Zea mays, Arabidopsis thaliana,* and the parasitic plant *Striga hermonthica. Striga hermonthica* was included as it is currently an unannotated species with no *MIRNAs* cataloged in miRBase, allowing us to test novel *MIRNA* validation. Each annotation tool was run using the same five libraries used for the plant species in the annotated *MIRNA* dataset (Table 2). For *S. hermonthica*, novel small RNA-seq libraries were generated from leaf and haustorial tissue. miRScore was then run using annotation results and the sRNA-seq data used for annotation.

miRDeep-P2 annotated the largest number of *MIRNA*s in each species, with over 2000 *MIRNA*s from *O. sativa*. The majority of them failed miRScore evaluation (Figure 4A, Supplemental File S3). Nearly all *MIRNA*s annotated by ShortStack passed miRScore inspection, (Figure 4B, Supplemental File S3). miRador annotations had a pass rate between 80 and 95% with failing loci flagged for various criteria (Supplemental File S3). For example, 168 failed because due to no miRNA/miRNA* reads detected, and 45 failed due to a lack of a 2nt 3’ overhang. Overall, miRScore effectively evaluated the outputs of several *MIRNA* annotation tools in plants, confirming its utility in diverse annotation workflows.

**Figure 4.**
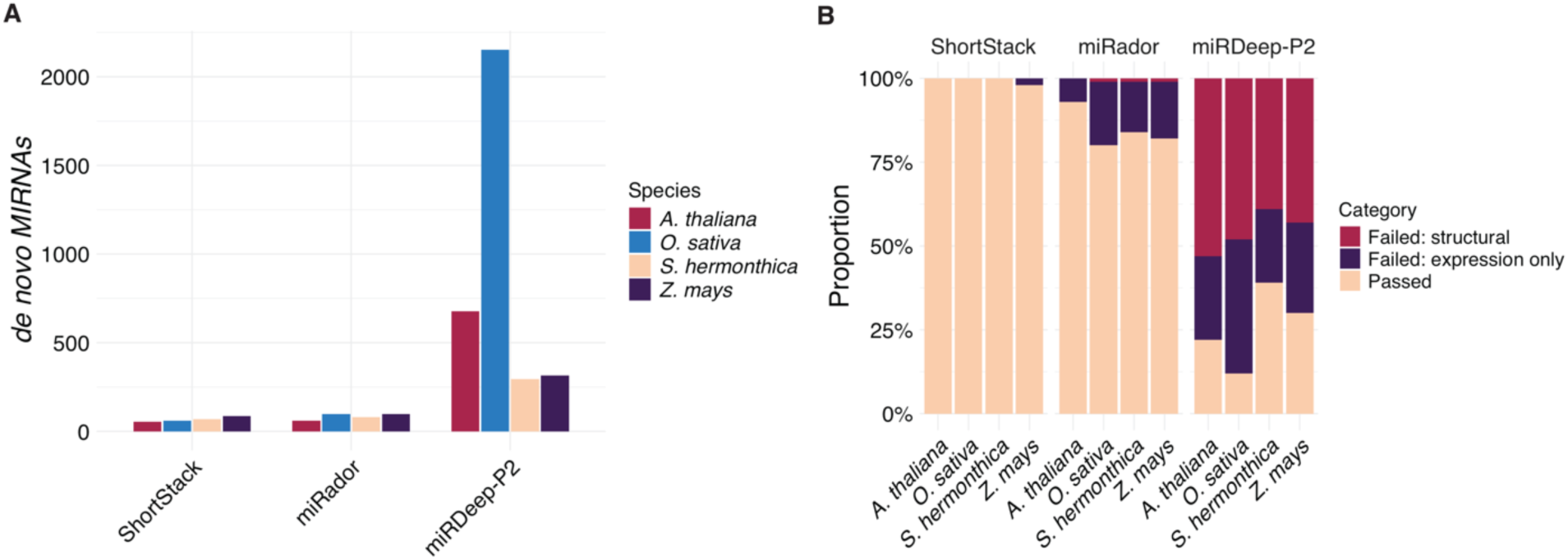
Evaluation of *de novo MIRNA* annotations. (A) The number of *MIRNAs* predicted by three *de novo* annotation programs by species. (B) Proportion of *de novo MIRNA*s which passed miRScore evaluation in each species.

## Availability and future directions

miRScore is an open-source python code and instructions are available on GitHub at https://github.com/Aez35/miRScore. A Bioconda recipe for miRScore is available, allowing easy installation using the conda package manager (30). In this study, we demonstrate that miRScore effectively validates *MIRNAs* in both annotated and novel *MIRNA* datasets across plant and animal species. miRScore enables rapid and robust analysis of *MIRNA* loci, without requiring a reference genome, and offers detailed metrics and visualizations for each locus to support comprehensive analysis of *MIRNA* datasets. We are confident that miRScore will contribute to improving *MIRNA* annotation quality and provide a tool for researchers to quickly verify annotations prior to downstream analysis. In addition, miRScore will provide an automated validation tool for miRNA databases and reduce the time it takes to catalog novel *MIRNAs.* miRScore’s ability to identify high confidence *MIRNAs* using widely accepted criteria quickly and accurately provides a valuable tool for researchers and database curators, contributing to the enhancement of *MIRNA* annotation quality in future studies.

## Methods

### Annotated datasets

Annotated *MIRNAs* from *Oryza sativa, Arabidopsis thaliana, Zea mays, Mus musculus,* and *Homo sapiens* were tested using miRScore version 0.2.0. Mature miRNA and hairpins sequences for each species were downloaded from miRBase version 22.1. The miRNA file submitted to miRScore for each species contained sequences for both the annotated miRNA and miRNA* sequence where applicable. Several miRNAs in each species began at position one of their respective hairpins and did not allow for evaluation of the miRNA duplex, particularly the two-nucleotide 3’ overhang structure. In these instances, miRScore will report the offending sequences and quit the run. These miRNAs were therefore removed from the dataset prior to running miRScore. sRNA-seq data accession numbers used for evaluation are listed in Table 2.

### Processing and alignment of small RNA-seq data

Small RNA-seq data for each dataset were trimmed to remove 3’ adapters using the ‘-autotrim’ feature of miRScore. Trim keys for each dataset were set to the miRScore default of ath-miR166a (UCGGACCAGGCUUCAUUCCCC) for plants and hsa-let7a (UGAGGUAGUAGGUUGUAUAGUU) for animals. The trimmed sRNA-seq data was aligned to hairpin sequences using bowtie (29) version 1.3.1. The BAM alignment files were merged and used to count reads that mapped to each hairpin using samtools (27) version 1.20.

### Striga hermonthica growth and sRNA-seq library preparation

*Striga hermonthica* kibos ecotype was grown on host *Oryza sativa* ssp. Japonica variety KitaakeX under 16-hour light conditions in a quarantine facility at 30°C for 45 days. Haustorium and leaf tissue were collected from *S. hermonthica* and total RNA was extracted using Zymo Quick-RNA Plant Miniprep Kit. sRNA-seq libraries were prepared essentially as described in Maguire et al., (2020) (40). Sequencing of the prepared libraries was performed on an Illumina NextSeq2000. New small RNA-seq libraries from *S. hermonthica* have been deposited at NCBI GEO under accession GSE282265.

### de novo MIRNA datasets

*MIRNAs* were annotated in four plant species using ShortStack version 4.0.4 (17), miRador commit c68c153 (19), and miRDeep-P2 version 1.1.4 (21). All annotation software were run on default settings for *de novo MIRNA* discovery. Genome assembly versions were: *Arabidopsis thaliana* (TAIR 10), *Oryza sativa* (IRGSP-1.0)*, Striga hermonthica* assembly SHERM (GCA_902706635.1)*, and Zea mays* (Zm-B73-REFERENCE-NAM-5.0). sRNA-seq data for *Striga hermonthica* was generated as described above. sRNA-seq data for the other plant species were acquired from accession numbers in Table 2.

Mature miRNA and hairpin sequences from resulting annotations were parsed and saved to two separate FASTA files. These FASTA files were used to test validation of *de novo MIRNAs* using miRScore version 0.2.0. The same sRNA-seq data used for *MIRNA* annotation were used to run miRScore for each species.

## Acknowledgments

This work was supported by an award from the US National Science Foundation to MJA (2130884), and a seed grant from The Huck Institutes of the Life Sciences at Penn State. We thank the Penn State Genomics Core Facility (RRID: SCR_023645) for sRNA-seq services.

## Author Contributions

AV wrote miRScore, performed testing and analysis, produced tables and figures, and drafted the manuscript. SGJ, BM, and MJA helped design the overall parameters of miRScore. SS supported small RNA-seq of *Striga hermonthica* samples. MJA tested miRScore and edited the manuscript.

## Disclosures

The authors declare no conflict of interest.

## Notes

### Competing Interest Statement

The authors have declared no competing interest.

