## Supplemental Figure S1 for "miRScore: a rapid and precise microRNA validation tool"

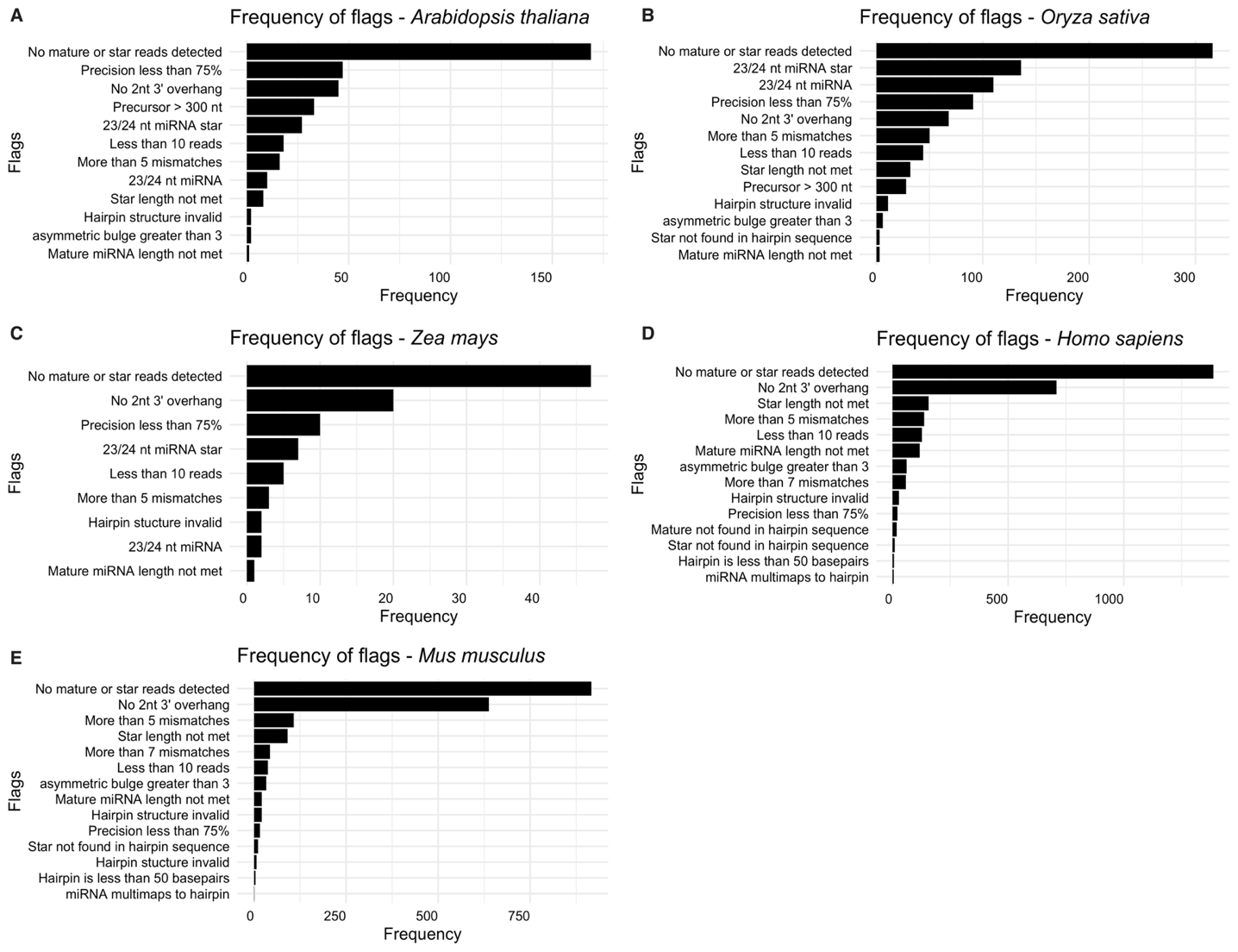

**Supplemental Figure 1.** Distribution of flags for all loci in annotated miRNA datasets. Some loci have multiple flags associated and are represented multiple times. (A) Frequency of miRScore flags in *A. thaliana.* (B) Frequency of miRScore flags in *O. sativa.* (C) Frequency of miRScore flags in *Z. mays.* (D) Frequency of miRScore flags in *H. sapiens*. (E) Frequency of flags in *M. musculus.*
